# Fluorescence analysis of serum lipoproteins using a histidine-imidazole polyacrylamide slab gel electrophoresis method

**DOI:** 10.1101/2025.04.26.650752

**Authors:** Yasuhiro Takenaka, Ikuo Inoue, Masaaki Ikeda, Yoshihiko Kakinuma

## Abstract

Polyacrylamide gel electrophoresis (PAGE) has traditionally been used to analyze lipoproteins in human and animal sera. Most conventional PAGE systems for lipoprotein analysis use disc electrophoresis, which limits the number of samples simultaneously analyzed and does not allow the comparison of electrophoretic profiles of multiple samples under the same conditions. Other methods, such as high performance liquid chromatography and agarose gel electrophoresis, are also used for lipoprotein analysis; however, they require specialized equipment and expertise. Here, we report an improved slab PAGE method with high-throughput, cost-effective, simple, rapid, and reproducible analysis of lipoproteins in the human serum. This system uses Tris-histidine as the running buffer and imidazole buffer for the polyacrylamide gel; therefore we named it the histidine-imidazole polyacrylamide gel electrophoresis (HI-PAGE) system. HI-PAGE effectively prevented band distortions of lipoproteins within an hour of electrophoresis and guaranteed precise quantitative analyses of multiple samples containing lipid-protein complexes. Furthermore, we pre-stained the lipoproteins with a fluorescent dye, Nile Red, and applied it to HI-PAGE electrophoresis. Fluorescent HI-PAGE (fHI-PAGE) was applied to the clinical samples and revealed that the method is highly sensitive and allows the quantitative detection of lipoproteins in human serum.

## 1 Introduction

Lipoproteins such as chylomicrons, very low-density lipoproteins (VLDL), intermediate-density lipoproteins (IDL), low-density lipoproteins (LDL), and high-density lipoproteins (HDL) are present in the human blood [1]. Among these, the cholesterol contained in LDL (LDL-c) is known as bad cholesterol. Cholesterol in IDL and small dense LDL is also called “super bad cholesterol” and causes atherosclerosis and myocardial infarction. Various analytical methods have been developed to predict the risk of arteriosclerosis and myocardial infarction by detecting IDL and small dense LDL. Among them, polyacrylamide gel electrophoresis (PAGE) [2–10], agarose gel electrophoresis [7,11–13], and chromatography [14,15] have been used since the 1960s to separate and analyze lipoproteins, and have many advantages such as low cost of equipment and reagents, easy handling, and short analysis time.

In PAGE, non-denaturing polyacrylamide disc gel analysis is frequently used to separate and evaluate serum lipoproteins. For disc electrophoresis of lipoproteins, a mixture of serum sample and staining dye was loaded onto a round glass tube filled with polyacrylamide and electrophoresis is performed [3,6,8,16,17]. This method is also used in general clinical tests. However, disc electrophoresis has limitations when comparing multiple samples under the same conditions, because only one sample can be analyzed per circular glass tube.

To solve this problem, we developed a slab-type PAGE method that can process multiple samples under the same conditions and is clinically applicable in terms of reproducibility, resolution, running time, and simple handling, by adding a number of technical improvements to conventional native PAGE methods.

## 2 Materials and methods

### 2.1 Reagents

Acrylamide, N,N’-ethylenebisacrylamide, imidazole, histidine, tris(hydroxymethyl)aminomethane (Tris), ammonium peroxodisulfate (APS), N,N,N’,N’-tetramethylethylenediamine (TEMED), Sudan Black B, Nile Red, and human serum albumin were purchased from FUJIFILM Wako Chemicals. β-BODIPY FL C12-HPC (D3792) was purchased from Thermo Fisher Scientific.

### 2.2 Study subject

Twenty-five individuals aged 40–85 years were recruited from the Department of Diabetes and Endocrinology at Saitama Medical University for this study registered in the University Hospital Medical Information Network Clinical Trial Registry (UMIN000040373 and UMIN000050974), a non-profit organization in Japan that meets the requirements of the International Committee of Medical Journal Editors. Eligible volunteers were those without a personal or family history of type 1 or 2 diabetes mellitus, pancreatitis, high blood pressure, angina pectoris, or coronary heart disease. All participants had fasting glucose levels below 110 mg/dL, body mass index below 32 kg/m^2^, total cholesterol (TC) < 220 mg/dL, triglycerides (TGs) < 200 mg/dL, smoked fewer than five cigarettes daily, and exhibited normal hepatic, thyroid, and renal functions, as determined by routine laboratory analyses. Clinical LDL-c level was calculated using the Friedewald equation:

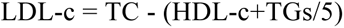

where TC, TG, and HDL-c values were measured at the central laboratory of Saitama Medical University Hospital. Informed consent was obtained from all the participants. This study was approval by the ethics committee of Saitama Medical University (20033.02 and 2023-021). Sera from healthy individuals (YT and YK) for the examination of electrophoresis conditions were collected with the approval of the Ethics Committee of Nippon Medical University (A-2021-057). Standard test serum (Cholestest^TM^ N Calibrator) was purchased from Sekisui Medical Co. Ltd. and prepared according to the manufacture’s instructions.

### 2.3 Isolation of HDL, LDL, and VLDL

HDL, LDL, and VLDL were sequentially isolated using the floating method [18,19]. Human serum from YT was mixed with an equal volume of 0.9% NaCl solution and centrifuged at 435,000 × *g* for 2.5 hours at 16°C using a TL-100 tabletop ultracentrifuge, Type 42.2 Ti (Beckman Coulter). Lipoproteins floating in the upper part of the tube were isolated as VLDL. The bottom portion of the tube (0.5 mL) was mixed with a 16.7% NaCl solution (0.5 mL) and centrifuged at 435,000 × *g* for 2.5 hours. The lipoprotein floating at the top was isolated as LDL, and the bottom portion of the tube (0.5 mL) was collected as HDL.

### 2.4 Acrylamide Gel Preparation of HI-PAGE

Non-gradient uniform acrylamide gels were casted onto conventional glass plates sealed with a gasket (Nihon Eido Co. Ltd.). We prepared a 30% acrylamide-bisacrylamide mixture (19:1) using N,N’-ethylenebisacrylamide to provide a large pore size and maintain physical gel strength (Table 1) [20]. The volume of each reagent used to prepare a single gel (90 × 70 × 1 mm, wide × height × thickness) is summarized in Table 2. First, upper and lower gel solutions, excluding TEMED, were prepared. An appropriate amount of TEMED was then added to the lower gel solution, thoroughly mixed, and poured onto the glass plates. Next, the lower gel solution was gently overlaid with distilled water and left to stand at 25°C for at least 10 min. After the lower gel solution solidified, the overlying distilled water was completely removed. The upper gel solution was mixed with TEMED, poured, and the comb was inserted. The mixture was then incubated at 25°C for more than an hour. To prevent over-drying, the gel was wrapped in plastic, and stored overnight in a refrigerator.

**Table 1.** Composition of 30% acrylamide-bisacrylamide mixture (19:1)

|  |  | Unit |
| --- | --- | --- |
| Acrylamide | 11.4 | g |
| N,N'-ethylenebisacrylamide | 0.6 | g |
| Ultrapure water | appropriate volume | mL |
| Total | 40 | mL |

**Table 2.** Composition of the upper and lower gels in HI-PAGE.

|  | Upper gel |  | Lower gel |  |
| --- | --- | --- | --- | --- |
|  | Volume (mL) | Concentration | Volume (mL) | Concentration |
| 1 M Tris-HCl, pH8.0 | 0.375 | 0.125M | - |  |
| 1 M imidazole-HCl, pH8.0 | - |  | 2.40 | 0.370 M |
| 30% acrylamide-bisacrylamide mixture (19:1) | 0.300 | 3.00% | 0.870 | 4.00% |
| Ultrapure water | 2.33 |  | 3.14 |  |
| 10% APS | 0.0380 |  | 0.0820 |  |
| TEMED | 0.00450 |  | 0.0100 |  |
| Total | 3.05 |  | 6.50 |  |

### 2.5 Running buffers for HI-PAGE

The preparation of the HI-PAGE running buffer is presented in Table 3. Weighed Tris and histidine were dissolved in an appropriate amount of ultrapure water, mixed thoroughly, then stored at 4°C. pH adjustment was not necessary; however, when the buffer was prepared as shown in Table 2, the pH was approximately 8.4. The amount of HI-PAGE running buffer to be prepared was adjusted according to the size of the electrophoresis tank.

**Table 3.** Composition of the HI-PAGE running buffer.

|  |  |  |
| --- | --- | --- |
|  |  | Final concentration |
| Tris | 0.91 g | 0.025 M |
| Histidine | 6.0 g | 0.13 M |
| Ultrapure water | up to 300 mL |  |

### 2.6 Preparation of the Pre-Staining Solution

Sudan Black B powder (0.3 g) was dissolve in 10 mL of dimethyl sulfoxide (DMSO) to prepare a 3% (w/v) Sudan Black DMSO solution, which was stored at room temperature. However, as staining properties gradually decrease, the solution should be prepared every 2–3 months. The pre-staining solution was prepared by mixing the components listed in Table 4 from top to bottom. It is important to follow this sequence, because altering it can result in incomplete dispersion of Sudan Black B throughout the solution, causing it to precipitate as insoluble granules. Each component was mixed vigorously upon addition. The pre-staining solution was stored in a refrigerator and prepared every 2–3 months because of the gradual decrease in its staining properties. For the fluorescent labeling of lipoproteins, NileRed, Tris buffer, and glycerol were mixed, as shown in Table 5.

**Table 4.** Composition of the pre-staining solution.

|  | Volume (μL) | Final concentration |
| --- | --- | --- |
| 3% Sudan Black DMSO solution | 40 | 0.12% |
| 10% DDM | 10 | 0.10% |
| 80% glycerol | 250 | 20% |
| 1 M Tris-HCl, pH7.4 | 50 | 0.05 M |
| Ultrapure water | 650 |  |
| Total | 1000 |  |

**Table 5.** Composition of the fluorescence pre-staining solution.

|  | Volume (μL) | Final concentration |
| --- | --- | --- |
| 50 mg/mL Nile Red DMSO solution | 50 | 2.5 mg/mL |
| 1% bromophenol blue | 37.5 | 0.0375% |
| 80% glycerol | 250 | 20% |
| 1 M Tris-HCl, pH7.4 | 50 | 0.05 M |
| Ultrapure water | 612.5 |  |
| Total | 1000 |  |

### 2.7 Sample Preparation

Before loading the samples onto the upper gel, the human serum was pre-stained using the solution listed in Table. 5. The ratio of the sample to the pre-staining solution was adjusted based on the concentration and total amount of lipoproteins in the sample. Human serum from healthy individual (YT) was used in this study (Fig. 1).

**Figure 1.**
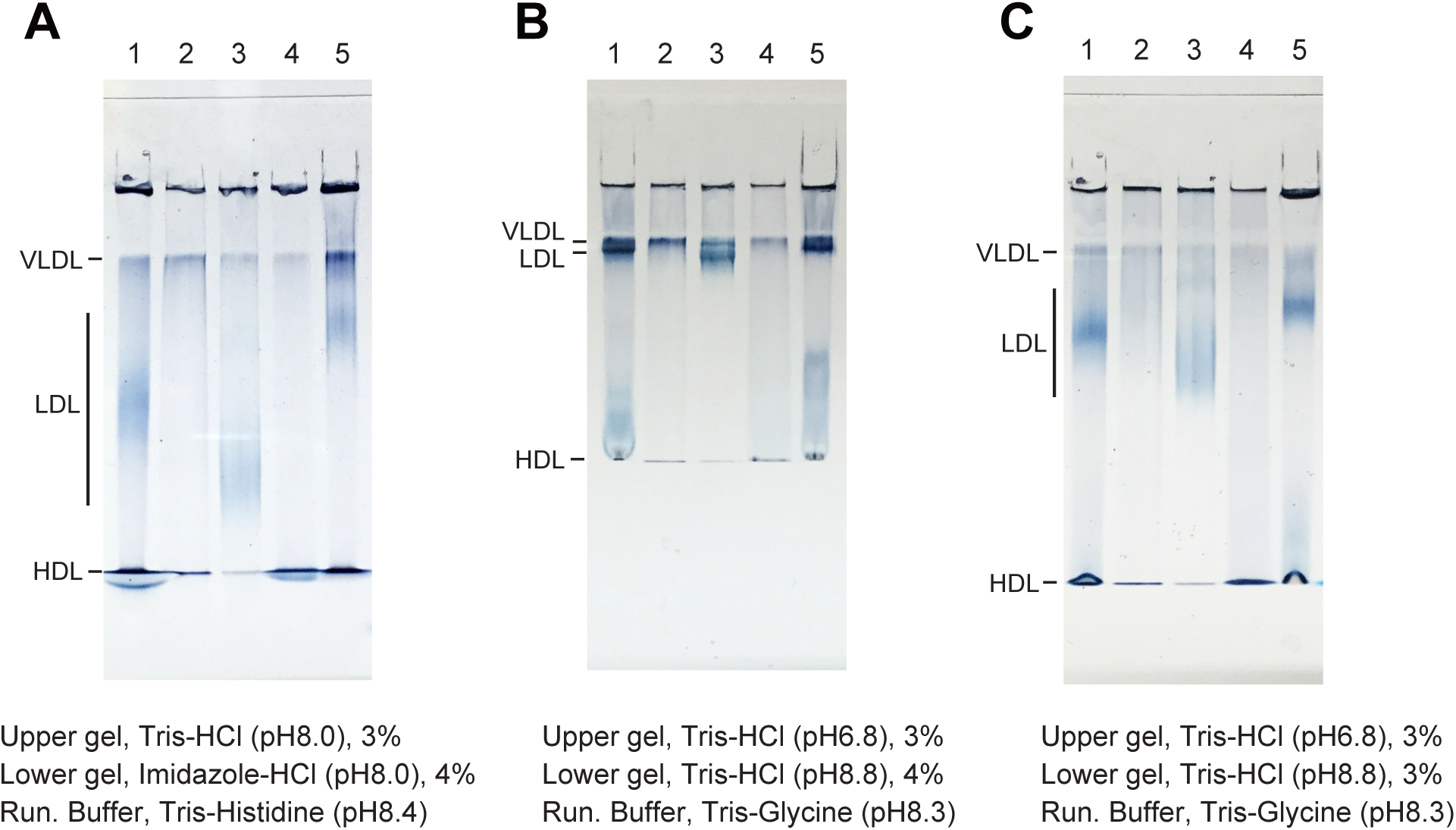
Electrophoresis of human lipoproteins using histidine-imidazole polyacrylamide gel electrophoresis (HI-PAGE). (A) Serum, VLDL, LDL, and HDL from healthy individual (lanes 1–4, respectively) and standard test serum (lane 5) were analyzed using colorimetric HI-PAGE. Compositions of the upper, lower, and running buffers are indicated below the gels. The positions of VLDL, LDL, and HDL are shown on the left side of the gel. (B) Electrophoretic profiles of conventional native PAGE with Tris-glycine running buffer. The percentage of lower gel was 4%. (C) The percentage of lower gel was 3%.

### 2.8 Sample Application and Electrophoresis Conditions

After mixing the sample with the pre-staining solution, the mixture was incubated at room temperature for 5 min to allow Sudan Black B to bind to lipoproteins in the sample. The entire sample volume was then loaded onto the upper gel. Loading was performed slowly and carefully using loading tips. Once the stained samples were loaded, electrophoresis was performed at a constant current (20–40 mA). The duration of electrophoresis depended on the size of the gel; a gel measuring 90 × 70 × 1 mm (w × h × t) was run for approximately 30–40 min at 40 mA. **Detection of Lipoproteins and Densitometric Quantification**After electrophoresis, gel images were obtained either by photographing the gel or by capturing the fluorescence of lipoproteins using a laser scanner, while the gel remained sandwiched between gel plates. Owing to the soft nature of the gel, handling it after its removal from the gel plates was challenging. To obtain fluorescence images of HI-PAGE, ChemiDoc Touch (Bio-Rad) or FLA 7000 (with excitation at 473 nm and filter O580; GE Healthcare) was used.

## 3 Results

### 3.1 Colorimetric Analysis of Human Lipoproteins Using HI-PAGE

HI-PAGE analysis of human serum from healthy individuals revealed that the use of Tris(hydroxymethyl)aminomethane hydrochloride as a buffer in the upper gel and imidazole hydrochloride in the lower gel, combined with Tris-histidine running buffer, effectively prevented band distortions of lipoproteins within an hour of electrophoresis (Fig. 1A). Thus, this method guaranteed precise analyses of multiple samples containing lipid-protein complexes, minimizing the occurrence of smiling and/or smearing features even within a shorter running time. In contrast, traditional non-denaturing polyacrylamide slab gel analysis using Tris buffer for the lower gel and Tris-glycine running buffer resulted in smiling or smearing band profiles of HDL (Figs. 1B and 1C). Moreover, band separation between VLDL and LDL using the traditional method was insufficient (Fig. 1B) compared with the HI-PAGE results (Fig. 1A), which required a longer electrophoresis time.

### 3.2 Fluorescence Analysis of Human Lipoproteins Using HI-PAGE

To further enhance the sensitivity of HI-PAGE, we pre-stained the standard test serum (CTN) with fluorescent dyes using Nile Red [21,22] or β-BODIPY FL C12-HPC (data not shown), as alternatives to Sudan Black B, before HI-PAGE electrophoresis. At the end of the run, the gel was immediately visualized using laser scanning or a fluorescent CCD imager, without separating the gel from the glass plates. As shown in Fig. 2A, the electrophoresis images of VLDL and LDL stained with Nile Red were comparable to those obtained by colorimetric HI-PAGE (Fig. 1A). We also obtained standard curves for VLDL (Fig. 2B) and LDL (Fig. 2C) by quantifying band intensities in the fluorescence HI-PAGE (fHI-PAGE) gel images. However, upon examining the fluorescence profile of purified human albumin, the most abundant protein in the serum, using fHI-PAGE, we observed a significant signal with mobility similar to that of HDL in the gel (Fig. 2D, a band indicated by an asterisk in lane 2), presumably because of bilirubin fluorescence that binds to human albumin [23]. The signal at the HDL position was not observed in the human albumin sample in the colorimetric HI-PAGE analysis (Fig. 2E, lane 2), indicating that the band observed in HI-PAGE (Figs. 1A-C and 2E, lane 1) was derived from HDL. We attempted to separate HDL and albumin bound to the fluorescent substance by increasing the acrylamide concentration (6% lower acrylamide concentration) without success (Fig. 2F). Therefore, to specifically detect HDL using fHI-PAGE, further improvements are needed, such as optimization in the fluorescent dye used or reduction of the fluorescent signal from the fluorescent substance. Nonetheless, the current fHI-PAGE method is suitable for qualitative and quantitative analysis of LDL and VLDL because no detectable signal was observed at the position corresponding to LDL and VLDL in the human albumin sample (Fig. 2D, lane 2). The fHI-PAGE method was applied to the analysis of clinical samples (Fig. 2G), resulting in lipoprotein profiles similar to those obtained by conventional disc electrophoresis (Fig. 2H).

**Figure 2.**
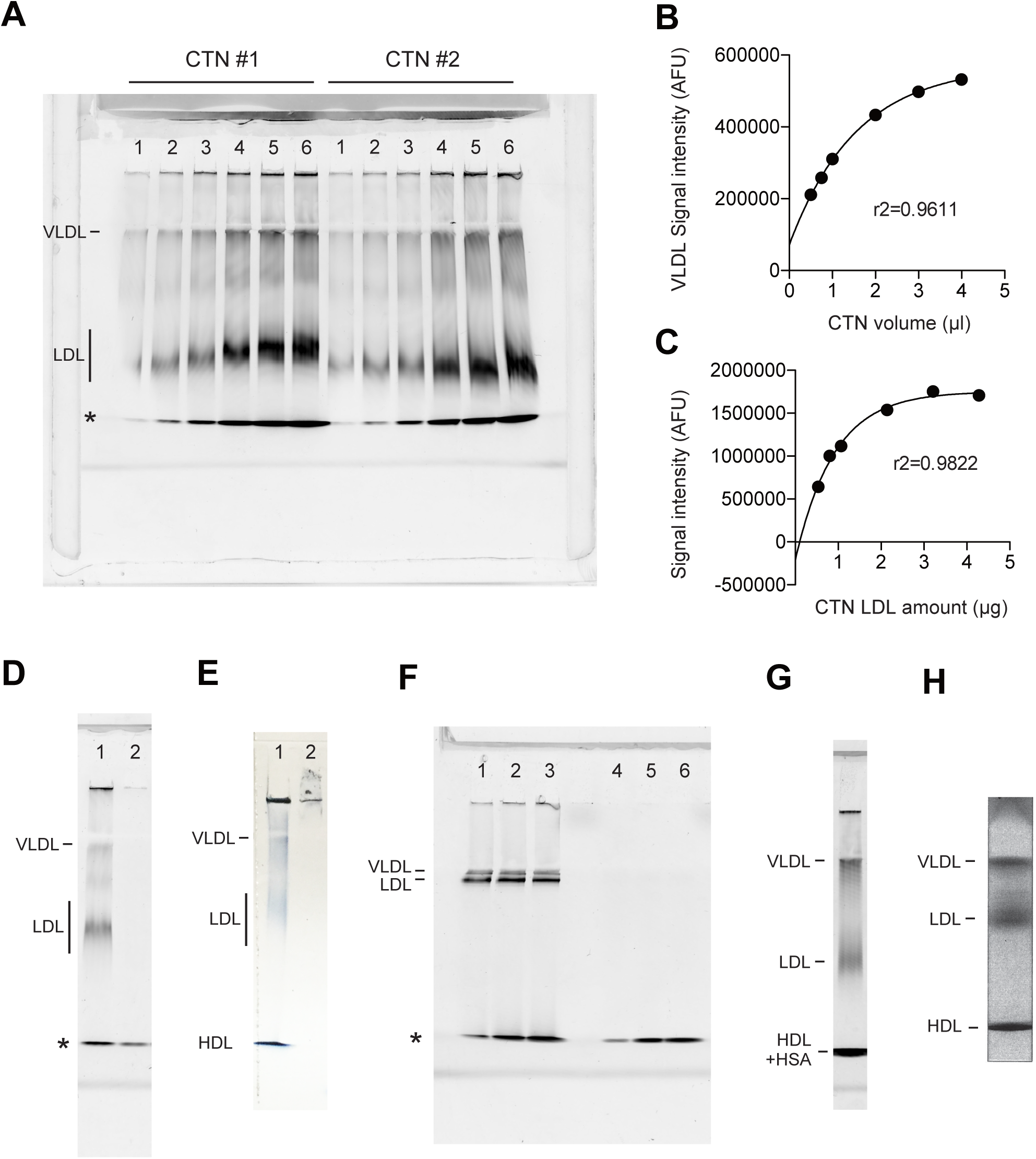
Electrophoresis of human lipoproteins using fluorescent HI-PAGE (fHI-PAGE). (A) Different amounts of standard test serum (CTN) were pre-stained with Nile Red and applied onto the gel (n = 2), and the fluorescent signal was imaged using laser scanning. Standard curves for VLDL (B) and LDL (C). (D) Human serum albumin (HSA) shows fluorescence at the size of HDL (asterisk). Lane 1, CTN; lane 2, human serum albumin. (E) Colorimetric HI-PAGE does not show a signal corresponding to the size of HDL. Lane 1, CTN; lane 2, human serum albumin. (F) fHI-PAGE analysis of different amounts of CTN (lanes 1-3) and HSA (lanes 5-7) with 6% acrylamide in the lower gel. An asterisk indicates the size of HDL. (G) fHI-PAGE analysis of the clinical sample. (H) Colorimetric disc electrophoresis of the same sample used in (G).

### 3.3 Fluorescence Analysis of LDL in Clinical Samples Using HI-PAGE

We applied fHI-PAGE to clinical samples from 18 patients to evaluate LDL-c levels. To calibrate LDL-c, we used varying amount of CTN along with patient serum samples for fHI-PAGE (Figs. 3A and 3B). Densitometry was performed on the LDL signal following laser scanning of the gel, and LDL-c was calculated using CTN standards. We then compared the LDL-c values obtained using fHI-PAGE with those calculated using the Friedewald equation (n = 22), and observed a moderate correlation between the two methods (R^2^ = 0.507) (Fig. 3C). To understand the discrepancies between these methods, we focused on several outlier samples. For example, the LDL-c values of patient B were 284 mg/dL and 78 mg/dL when calculated using the fHI-PAGE and Friedewald equation, respectively (Figs. 3C and 3D). This discrepancy is likely due to Patient B’s high triglyceride level (328 mg/dL), which led to a significantly lower LDL-c value when calculated using the Friedewald equation. In contrast, the LDL-c value for Patient G calculated using the Friedewald equation was 250 mg/dL, which was notably higher than that obtained using fHI-PAGE analysis (198 mg/dL). This may be attributed to the high total cholesterol level of Patient G (340 mg/dL). On the other hand, LDL-c could not be calculated using the Friedewald equation for Patient J because of an abnormally high triglyceride level (2,048 mg/dL), whereas the LDL-c level was successfully measured using fHI-PAGE analysis (Fig. 3D).

**Figure 3.**
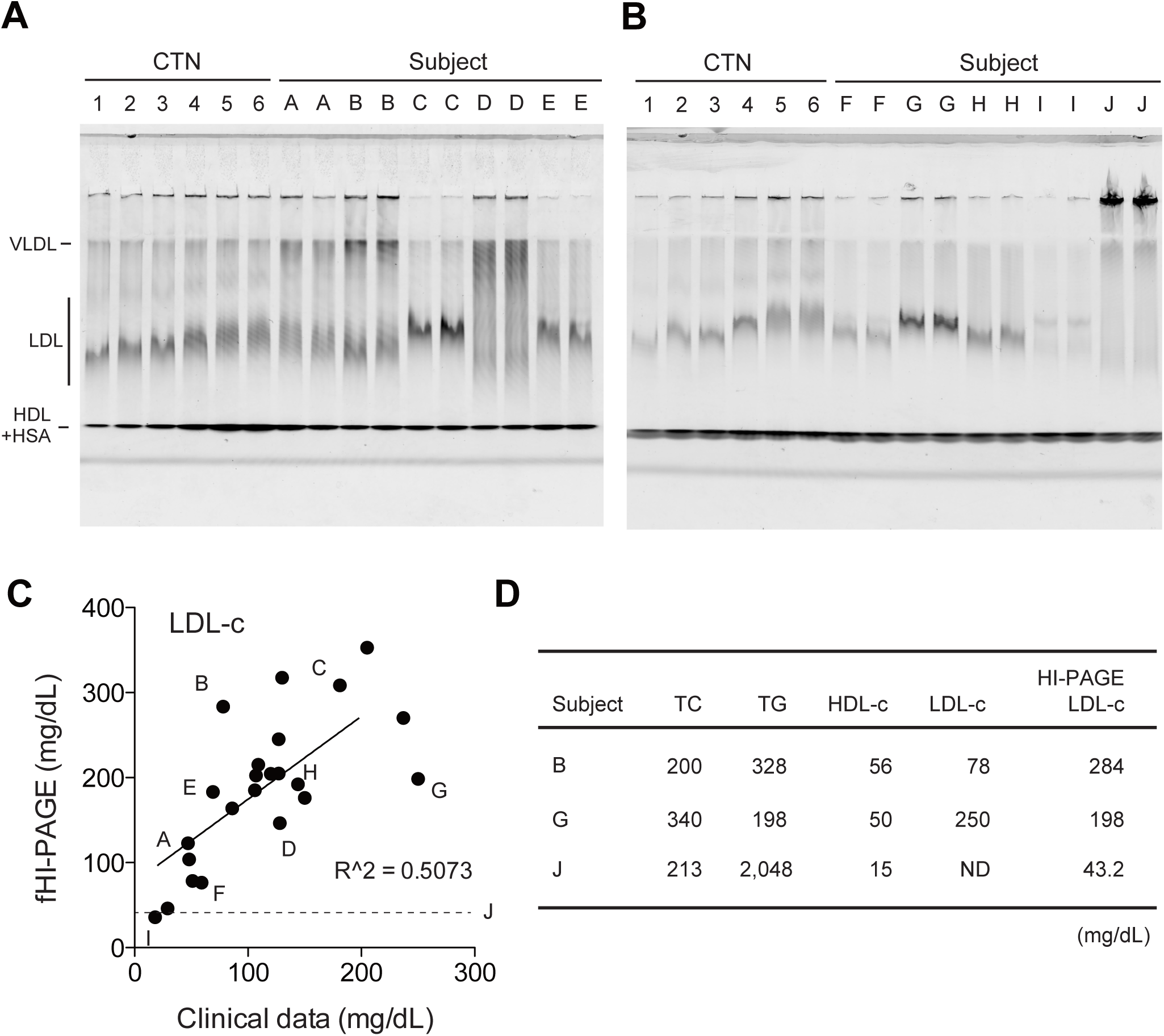
fHI-PAGE analysis of clinical samples. (A) CTN (lanes 1–6) and clinical samples (lanes A-E, duplicates) were analyzed using fHI-PAGE. (B) Additional gel image of fHI-PAGE with CTN (lanes 1–6) and clinical samples (lanes F-J, duplicates). (C) Dot plot of values obtained using densitometry analysis of the LDL signal of fHI-PAGE (*y* axis) and that calculated using the Friedewald equation (*x* axis). The letters correspond to those in Figs. 3A and 3B. (D) Total cholesterol (TC), triglyceride (TG), HDL-c, and calculated LDL-c values with fHI-PAGE LDL-c results of the outfitter samples. The unit of each value is mg/mL.

## 4 Discussion

We developed a rapid, simple, sensitive, and high-throughput slab-type PAGE system for lipoprotein analysis. This technology is based on the native PAGE method for lipoproteins, including conventional disc gel electrophoresis, with several modifications, such as buffer systems and dyes, to stain lipoproteins. Compared to disc electrophoresis, slab electrophoresis allows for a simpler operation as it eliminates the need to prepare multiple tube gels for handling multiple samples. In addition, it enables sample comparison under identical conditions on the same gel, which is advantageous for detecting subtle differences between the samples. Furthermore, because there were multiple wells, we obtained a calibration curve by preparing standard serum at several amounts, ranging from the upper to lower limits. In addition, whereas conventional disc PAGE requires 25 µL of serum for analysis, 5 and 1 µL are sufficient for HI-PAGE and fHI-PAGE analyses, respectively. Conventional disc PAGE requires photopolymerization of the loading gel for 30–45 min before electrophoresis, whereas our method allows immediate application after mixing serum and staining dye.

In conventional slab-type PAGE, the bands of each lipoprotein are prone to distortion owing to smearing or smiling during electrophoresis. Here, we improved the distortion of the band shapes of VLDL, LDL, and HDL by using imidazole as the lower gel buffer and Tris-histidine as the running buffer. Furthermore, the slab-type PAGE reported by Singh *et al*. requires a total run time of 190 min [2], whereas in the HI-PAGE system, electrophoresis can be completed within 40 min, which is almost the same as that of conventional disc PAGE.

Our method had some limitations: 1) limited linearity when the signal intensities of lipoproteins were scanned using a densitometer, especially at a higher amount of LDL; 2) a notable mobility shift of LDL was observed when increasing the loading amount, which caused difficulties in analyzing the slight mobility change of LDL between samples with different loading amounts; and 3) as shown in Fig. 2D, HDL was not measured accurately using fHI-PAGE. Nevertheless, this method has numerous advantages, including affordability, simplicity, and the ability to perform real-time monitoring during analysis without expensive equipments such as high performance liquid chromatography [24] or other specialized analytical techniques. We believe that further improvements and applications of this method have a significant potential.

## 5 Concluding remarks

In conclusion, we developed a colorimetric and fluorescent slab PAGE method with high-throughput, cost-effective, simple, rapid, and reproducible analysis of lipoproteins in the human serum using Tris-histidine as the running buffer and imidazole buffer for the polyacrylamide gel (HI-PAGE). Fluorescent HI-PAGE was applied to the clinical samples and revealed that the method is highly sensitive and allows the quantitative detection of lipoproteins in human serum.

## Data availability statement

Data supporting the findings of this study are available from the corresponding author upon reasonable request.

## Acknowledgments

The authors are grateful to Ms. Sawako Sato and Ms. Megumi Kumagai for technical assistance. We acknowledge ChatGPT (GPT-4o), developed by OpenAI and Editage (www.editage.jp) for their invaluable assistance in editing the English text.

## Author contributions

Y.T., I.I., M.I.: conceptualization, data curation, investigation, methodology, visualization, and writing - original draft. Y.K.: supervision of the sample preparation and writing - original draft.

## References

[1] Feingold KR. Introduction to Lipids and Lipoproteins. In: Feingold KR, Anawalt B, Blackman MR, Boyce A, Chrousos G, Corpas E, et al., editors. Endotext. South Dartmouth (MA): MDText.com, Inc.

[2] Singh Y, Lakshmy R, Gupta R, Kranthi V. A rapid 3% polyacrylamide slab gel electrophoresis method for high through put screening of LDL phenotype. Lipids Health Dis. 2008;7:47.

[3] Hoefner DM, Hodel SD, O’Brien JF, Branum EL, Sun D, Meissner I, et al. Development of a rapid, quantitative method for LDL subfractionation with use of the Quantimetrix Lipoprint LDL System. Clin Chem. 2001;47:266–74.

[4] Fonda M, Semolic AM, Soranzo MR, Cattin L. Production of polyacrylamide gradient gel for lipoprotein electrophoretic separation. Clin Chim Acta. 2003;338:73–8.

[5] Krauss RM, Burke DJ. Identification of multiple subclasses of plasma low density lipoproteins in normal humans. J Lipid Res. 1982;23:97–104.

[6] Narayan KA, Creinin HL, Kummerow FA. Disc electrophoresis of rat plasma lipoproteins. J Lipid Res. 1966;7:150–7.

[7] Inoue I, Koh HS, Mizotani K, Goto S, Tanaka K, Yagasaki F, et al. A patient with severe hypertriglyceridemia associated with anemia and hypoalbuminemia. J Atheroscler Thromb. 2003;10:192–201.

[8] Bando Y, Tohyama H, Aoki K, Kanehara H, Hisada A, Okafuji K, et al. Ipragliflozin lowers small, dense low-density lipoprotein cholesterol levels in Japanese patients with type 2 diabetes mellitus. Journal of clinical & translational endocrinology. 2016;6:1–7.

[9] Koizumi T, Kaneda H, Komiyama N, Inoue I, Muramatsu T, Nakajima K. Lipoprotein Profiles before Heparin Administration in Patients with or without Coronary Thrombosis Following Atherosclerosis. Annals of vascular diseases. 2021;14:31–8.

[10] Davis BJ. DISC ELECTROPHORESIS. II. METHOD AND APPLICATION TO HUMAN SERUM PROTEINS. Ann N Y Acad Sci. 1964;121:404–27.

[11] Freeman LA, Shamburek RD, Sampson ML, Neufeld EB, Sato M, Karathanasis SK, et al. Plasma lipoprotein-X quantification on filipin-stained gels: monitoring recombinant LCAT treatment ex vivo. J Lipid Res. 2019;60:1050–7.

[12] Burstein M, Scholnick HR, Morfin R. Rapid method for the isolation of lipoproteins from human serum by precipitation with polyanions. J Lipid Res. 1970;11:583–95.

[13] Kido T, Kurata H, Matsumoto A, Tobiyama R, Musha T, Hayashi K, et al. Lipoprotein analysis using agarose gel electrophoresis and differential staining of lipids. J Atheroscler Thromb. 2001;8:7–13.

[14] Chopra M, Fitzsimons P, Hopkins M, Thurnham DI. Dialysis and gel filtration of isolated low density lipoproteins do not cause a significant loss of low density lipoprotein tocopherol and carotenoid concentration. Lipids. 2001;36:205–9.

[15] Vedie B, Myara I, Pech MA, Maziere JC, Maziere C, Caprani A, et al. Fractionation of charge-modified low density lipoproteins by fast protein liquid chromatography. J Lipid Res. 1991;32:1359–69.

[16] Matsuo K, Inoue I, Matsuda T, Arai T, Nakano S. Relative increase in production ratio of small dense low-density lipoprotein in acute coronary syndrome with high coronary plaque burden: an ex-vivo analysis. Heart Vessels. 2024.

[17] Nakano T, Inoue I, Seo M, Takahashi S, Awata T, Komoda T, et al. Rapid and simple profiling of lipoproteins by polyacrylamide-gel disc electrophoresis to determine the heterogeneity of low-density lipoproteins (LDLs) including small, dense LDL. Recent Pat Cardiovasc Drug Discov. 2009;4:31–6.

[18] Cole TG, Ferguson CA, Gibson DW, Nowatzke WL. Optimization of beta-quantification methods for high-throughput applications. Clin Chem. 2001;47:712–21.

[19] Schumaker VN, Puppione DL. Sequential flotation ultracentrifugation. Methods Enzymol. 1986;128:155–70.

[20] Hochstrasser DF, Patchornik A, Merril CR. Development of polyacrylamide gels that improve the separation of proteins and their detection by silver staining. Anal Biochem. 1988;173:412–23.

[21] Greenspan P, Gutman RL. Detection by nile red of agarose gel electrophoresed native and modified low density lipoprotein. Electrophoresis. 1993;14:65–8.

[22] Greenspan P, Lou P. Spectrofluorometric studies of nile red treated native and oxidized low density lipoprotein. Int J Biochem. 1993;25:987–91.

[23] Jacobsen J. Binding of bilirubin to human serum albumin - determination of the dissociation constants. FEBS Lett. 1969;5:112–4.

[24] Yamashita S, Okazaki M, Okada T, Masuda D, Yokote K, Arai H, et al. Distinct Differences in Lipoprotein Particle Number Evaluation between GP-HPLC and NMR: Analysis in Dyslipidemic Patients Administered a Selective PPARα Modulator, Pemafibrate. J Atheroscler Thromb. 2021;28:974–96.

